# Confirming the Pro-longevity Effects of H3K4me3-deficient *set-2* Mutants in Extending Lifespan in *C. elegans*

**DOI:** 10.1101/2022.08.02.502497

**Authors:** Carlos G. Silva-García, William B. Mair

## Abstract

The COMPASS chromatin complex, which trimethylates lysine 4 on histone H3 (H3K4me3), regulates lifespan in *Caenorhabditis elegans*. Knockdown or partial loss-of-function of SET-2, a member of the COMPASS complex, extends the lifespan of worms. However, recent observations suggested that full loss of SET-2 methyltransferase activity via deletion of its active site reduces lifespan, indicating the degree of COMPASS inhibition may be critical to its effects on aging. To further explore these inconsistencies, we examined *set-2* longevity across a range of interventions from weak to full inhibition. Via CRISPR/Cas9 genome-editing, we made two new *set-2* mutants, a new genocopy of the most commonly used *set-2(ok952)* allele for mild *set-2* inhibition, and a full *set-2* genomic deletion. We found that both new strains (partial and null) and RNAi show a lifespan extension in *C. elegans* fed HT115 *E. coli*. However, neither mutant was long-lived in *C. elegans* fed OP50-1 *E. coli*. These data confirm that the previous lifespan extension observed in *set-2(ok952)* mutants was indeed the result of *set-2* inhibition (and not a secondary linked mutation generated in the original strain). These data also indicate that a diet-dependent mechanism might contribute to the regulation of lifespan under H3K4me3 deficiency and highlight how COMPASS-mediated longevity involves a complex interaction between chromatin state and environment.

## MAIN TEXT

The eukaryotic, multicellular organism *Caenorhabditis elegans* has been well established as a model system to recapitulate many human diseases, and in recent decades it has become invaluable for experimental research at both the physiological and genetic levels *in vivo*. Since the first *C. elegans* mutants were generated in the 1970s, hundreds of novel mutants have been identified with relevance for understanding human disorders. However, traditional genetic methods to generate single strains sometimes carry other non-desired mutations that can influence *C. elegans* physiology (Kutscher and Shaham, 2014). The development of the CRISPR/Cas9 system now allows precise and rapid genome editing in the worm genome (Friedland et al., 2013) and has helped to circumnavigate this issue.

*C. elegans* has homologs of approximately two-thirds of all human disease genes and has been used to study metabolic and neurodegenerative disorders. In addition, due to its physiological characteristics and short lifespan, *C. elegans* is a powerful model for studies on aging and age-related diseases (Denzel et al., 2018). Since the discovery for the first time that a single mutation can extend lifespan in animals, many other *C. elegans* longevity mutants have been isolated (Kenyon, 2010). In addition to genetic modifications, epigenetic modifications and mutations in components that modulate chromatin have emerged as critical regulators of aging. In particular, partial loss-of-function mutations or knockdowns of members (SET-2, WDR-5, and ASH-2) of the complex of proteins associated with Set1 (COMPASS), which catalyzes trimethylation of lysine 4 on histone H3 (H3K4me3), extend lifespan in *C. elegans* (Greer et al., 2010, 2011; Han et al., 2017; Lee et al., 2019). However, recent data challenged this view, showing that partial mutant animals in the *set-2* gene have a reduced lifespan (Caron et al., 2021) and suggesting previous longevity effects may have been the indirect result of linked mutations generated during the construction of the original strains. Here, we used CRISPR/Cas9 and RNA interference (RNAi) to understand and reconcile the differences in these observations.

First, we knocked down *set-2* by RNAi in wild-type worms. As previously reported (Greer et al., 2010, 2011; Han et al., 2017), we observed a lifespan extension under *set-2* RNAi conditions (**Fig. 1a**). Then, using CRISPR/Cas9 genome-editing, we further evaluated reported inconsistencies in the role of *set-2* mutants in longevity. The most used partial loss-of-function allele of *set-2* is *ok952*. The *set-2(ok952)* strain was generated by trimethylpsoralen, which produces small deletions with an average size of one to three kilobases; however, all base transitions and transversions have been observed (Kutscher and Shaham, 2014). Despite multiple outcrosses, *set-2(ok952)* animals could possibly carry tightly linked mutations outside the *set-2* open reading frame that could influence animals’ physiology. Therefore, we decided to genocopy the original (*ok952)* allele and test its role in longevity. Using CRISPR, we generated exactly the same *ok952* deletion/insertion present in the *set-2* locus, *set-2(wbm94)*: 1269 base pairs deletion and 12 base pair insertion (**Fig. 1b**). Using previously reported conditions (Greer et al., 2010, 2011; Han et al., 2017), we found that *set-2(wbm94)* genocopy mutant animals have a lifespan extension phenotype similar to that observed in *set-2(ok952)* mutants (**Fig. 1d**).

**Figure 1.**
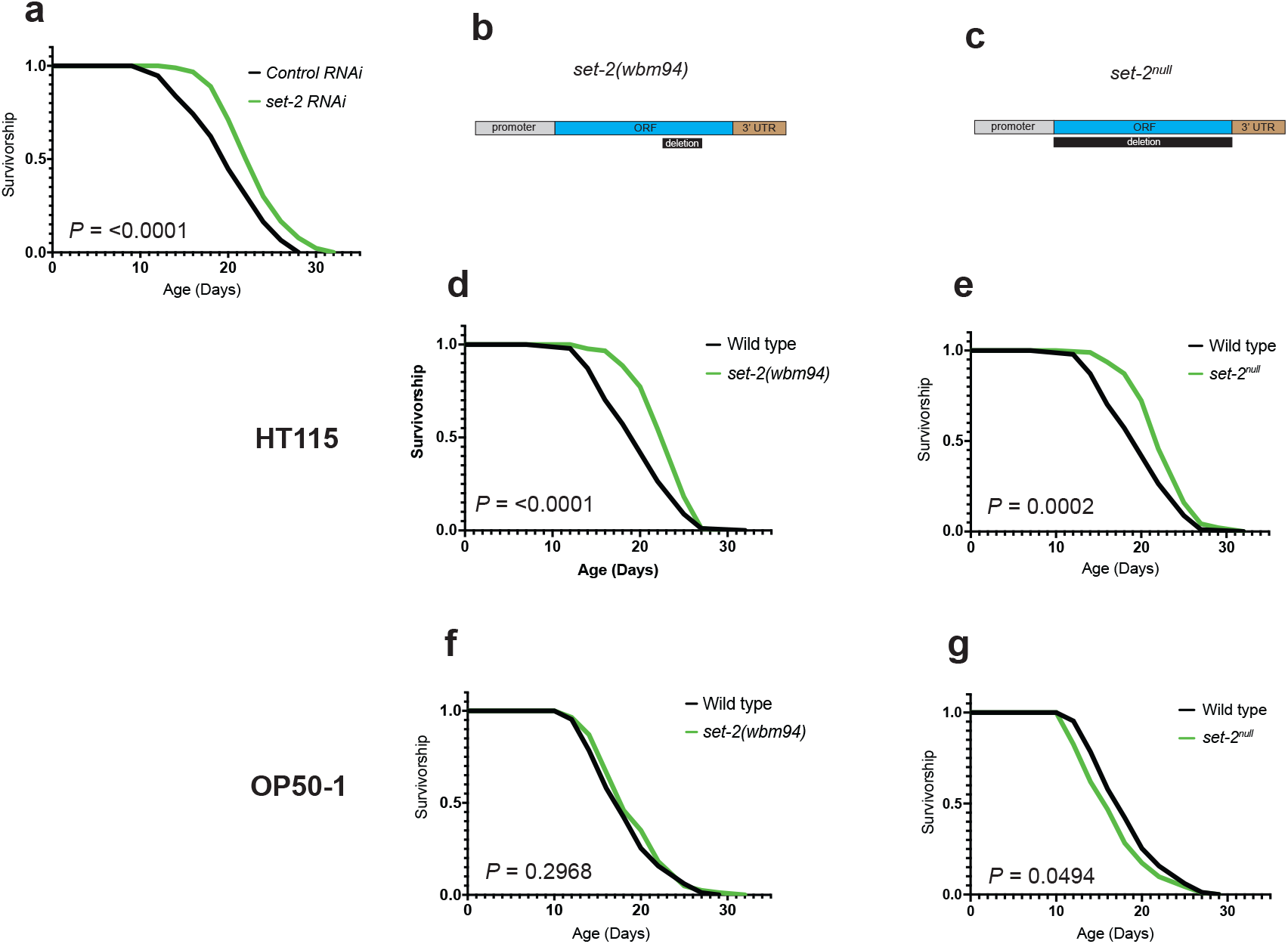
The lifespan trajectory of H3K4me3-deficient animals depends on the food source. **a)** Lifespan curves show that the knockdown of *set-2* extends the lifespan of wild-type animals. **b)** Schematic of the *set-2 wbm94* partial deletion, *set-2(wbm94)*. **c)** Schematic of the *set-2* full deletion, *set-2*^*null*^. **d)** *set-2(wbm94)* mutants show a lifespan extension on HT115. **e)** Lifespan curves showing that *set-2*^*null*^ mutants live longer on HT115. **f-g)** Lifespan curves showing that *set-2(wbm94)* and *set-2*^*null*^ mutants do not have a longevity phenotype on OP50-1. The Log-rank (Mantel-Cox) method was used to compare survival curves. *P* values inset in figure.

Next, we examined whether weak inhibition of SET-2 might be pro-longevity while strong inhibition might accelerate aging, as previously suggested (Caron et al., 2021). Since *set-2(wbm94)* and the original *set-2(ok952)* strain carry a partial loss-of-function of the SET-2 protein, we generated a full *set-2* null mutant. We used CRISPR to delete 7130 base pairs of the *set-2* gene to remove the entire coding sequence from START to STOP codons, *set-2*^*null*^ (**Fig. 1c**). We then performed a lifespan assay and found that *set-2*^*null*^ mutant animals lived long (**Fig. 1e**). Taken together, these data indicate that partial inhibition of SET-2, via RNAi knockdown [*set-2RNAi*] and partial loss-of-function mutation [*set-29(wbm94)*], or the complete loss-of-function [*set-2*^*null*^] of *set-2* all induce lifespan extension (**Fig. 1a-e**).

In an attempt to reconcile our results with previous work showing that point mutation in the catalytic domain (SET domain) of SET-2 reduces *C. elegans* lifespan (Caron et al., 2021), we compared the experimental conditions. A critical difference between studies was the husbandry. While the majority of previous work performed all lifespans on HT115 bacteria (Greer et al., 2010, 2011; Han et al., 2017), Caron *et al*. carried out the experiments on OP50-1 bacteria (Caron et al., 2021). Therefore, we decided to repeat the lifespan assays of our new strains on OP50-1. We found that neither *set-2(wbm94)* nor *set-2*^*null*^ mutants were long-lived on this food (**Fig. 1f-g**), similar to previous observations (Caron et al., 2021). These data indicate that bacterial diets can indeed alter lifespan trajectories in *set-2* mutant animals and suggest gene by environment interactions can underlie differences in published observations.

In summary, our data suggest there can be contextual lifespan extension via COMPASS-mediated deficiency of H3K4me3 in *C. elegans*, depending upon the food source. Diet is probably one of the most variable factors in life due to the range of options that organisms are exposed to in nature. *C. elegans* is a classic example of a model organism that has been disconnected from its natural ecology. The bacteria used in lab husbandry is not one they are exposed to in the wild (Samuel et al., 2016). Bacterial diets play essential roles in *C. elegans* physiology (Macneil and Walhout, 2013). Indeed, worms are able to respond to different bacterial exposures, altering physiology, lipid content, and lifespan across different diets (Brooks et al., 2009; Kim, 2013; Pang and Curran, 2014; Samuel et al., 2016; Stuhr and Curran, 2020). Beyond a food source, *E. coli* also represents a live microbial environment in which *C. elegans* live. Therefore, these types of contextual interactions between the genome of the host and associated microbes may also mimic the complex relationship between host genetics and the microbiome seen in higher organisms (Heintz and Mair, 2014). More work is necessary to understand the mechanism underlying this fascinating interaction between COMPASS activity and environment, specifically as it relates to lifespan regulation under a deficiency of H3K4me3.

## METHODS

### *C. elegans* strains and husbandry

N2 wild-wild type *C. elegans* strain was obtained from the Caenorhabditis Genetic Center, which is funded by the NIH Office of Research Infrastructure Programs (P40 OD010440). N2, *set-2(wbm95)* [*set-2*^null^], and *set-2(wbm94)* strains were maintained on standard nematode growth media (NGM) seeded with *E. coli* OP50-1 and maintained at 20 °C.

### Microbe strains

OP50-1 bacteria were cultured overnight in LB at 37 °C, after which 100 µl of liquid culture was seeded on plates to grow for two days at room temperature. Worms were grown at 20 °C on the *E. coli* OP50-1 or HT115 (empty vector, EV). RNAi experiments employed HT115 bacteria from the Ahringer library (Source Bioscience) expressing dsRNA against the gene noted or empty vector control. HT115 bacteria were cultured overnight in LB containing 100 mg ml^-1^ carbenicillin and 12.5 ml ml^-1^ tetracycline at 37 °C, after which 100 µl of LB was seeded on NGM plates containing 100 mg ml^-1^ carbenicillin to grow for two days at room temperature. dsRNA expression was induced by adding 100 µl IPTG (100 mM) at least 2 hours before worms were introduced to the plates.

### Microinjection and CRISPR/Cas9-triggered homologous recombination

All CRISPR edits and insertions required to generate the strains were performed using the previously described CRISPR protocol (Paix et al., 2015; Silva-García et al., 2019). Briefly, homology repair templates were amplified by PCR, using primers that introduced a minimum stretch of 35 bp homology at both ends. Single-stranded oligo donors (ssODN) were also used as repair templates. CRISPR injection mix reagents were added in the following order: 0.375 µl Hepes pH 7.4 (200 mM), 0.25 µl KCl (1 M), 2.5 µl tracrRNA (4 µg/µl), 0.6 µl *dpy-10* crRNA (2.6 μg/μl), 0.25 μl *dpy-10* ssODN (500 ng/μl), and PCR or ssODN repair template(s) up to 500 ng/µl final in the mix. Water was added to reach a final volume of 8 µl. 2 µl purified Cas9 (12 μg/μl) added at the end, mixed by pipetting, spun for 2 min at 13000 rpm, and incubated at 37 °C for 10 min. Using standard methods, mixes were microinjected into the germline of day one adult hermaphrodite worms (Evans, 2006).

### crRNAs and repair templates

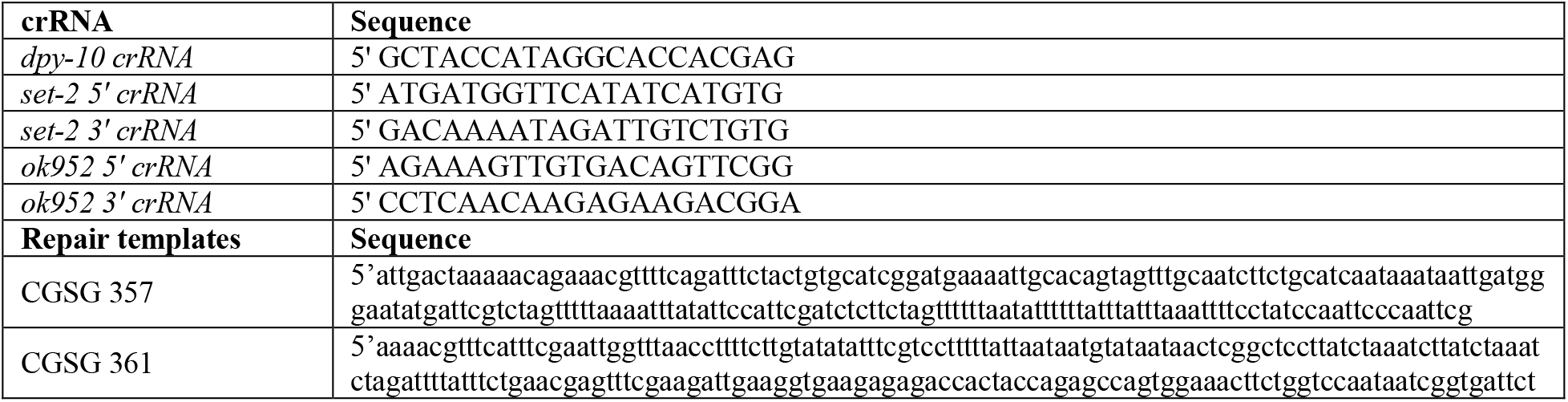

#### Survival Analyses

Lifespan experiments were performed as described previously (Burkewitz et al., 2015; Weir et al., 2017). Experiments were performed on standard NGM on OP50-1 or standard NGM containing 100 mg ml^-1^ carbenicillin at 20 °C and on HT115 (empty vector, EV). RNAi experiments were performed from hatching on standard NGM containing carbenicillin (100 mg ml^-1^). Expression of dsRNA was induced by pipetting 100 µl IPTG solution (100 mM, containing 100 mg ml^-1^ carbenicillin and 12.5 ml ml^-1^ tetracycline) onto HT115 lawns prior to placing worms. Worms were synchronized by bleaching using gravid adults. After bleaching, embryos were placed on plates. When the progeny reached adulthood (∼72 h), 100 worms were transferred to fresh plates with 20 worms per plate. Worms were transferred to fresh plates every other day until reproduction had ceased (day 9-12). Survival was scored every 1–2 days, and a worm was deemed dead when unresponsive to 3 taps on the head and tail. Worms were censored due to contamination on the plate, leaving the NGM, eggs hatching inside the adult, or loss of vulval integrity during reproduction. Lifespan analysis was performed using GraphPad Prism. The Log-rank (Mantel-Cox) method was used to compare survival curves.

